# Enrichment of allelic editing outcomes by Prime Editing in iPS cells

**DOI:** 10.1101/2024.03.19.585665

**Authors:** Ryo Niwa, Tomoko Matsumoto, Alexander Y. Liu, Maki Kawato, Takayuki Kondo, Kayoko Tsukita, Haruhisa Inoue, Thomas L. Maurissen, Knut Woltjen

**Author notes:** Correspondence should be addressed to K.W., Center for iPS Cell Research and Application (CiRA), Kyoto University, 53 Kawahara-cho, Shogoin, Sakyo-ku, Kyoto 606-8507, JAPAN.

## Abstract

Gene editing in human induced pluripotent stem (iPS) cells with programmable nucleases facilitates reliable disease models, but methods using double-strand break repair often produce random on-target by-products. Prime editing (PE) combines Cas9 nickase with reverse transcriptase (RT) and a prime editing guide RNA (pegRNA) encoding a repair template to reduce by-products. We implemented a GMP-compatible protocol for transfecting Cas9- or PE-2A-mCherry plasmids to track and fractionate human iPS cells based on PE expression level. We compared the editing outcomes of Cas9- and PE-based methods in a GFP-to-BFP conversion assay, at the *HEK3* benchmark locus, and at the *APOE* Alzheimer’s risk locus, revealing superior precision of PE at high expression levels. Moreover, sorting cells for PE expression level influenced allelic editing outcomes at the target loci. We expect that our findings will aid in the creation of gene-edited human iPS cells with intentional heterozygous and homozygous genotypes.

**Highlights:**

1. Delivered large plasmids to human iPS cells under GMP-compliant conditions
2. Developed a flow cytometry-based approach to enrich for PE in human iPS cells
3. Demonstrated few on-target indels in cells regardless of PE expression
4. Sorted iPS cells based on PE expression level to influence mono- or bi-allelic editing

## Introduction

Human induced pluripotent stem cells (iPS cells) are produced from somatic cells and have indefinite proliferative and differentiation potential^1–3^. In addition, they can differentiate into multiple cell types, while retaining their normal diploid karyotypes and genome from donors. Based on these characteristics, iPS cells are widely used as cellular-level models to study human genetic diseases. Genetic disease modeling is generally achieved by correcting mutations in iPS cells derived from patients with hereditary diseases, or by introducing mutations into iPS cells prepared from healthy donors without diseases^4^. In addition, modifying a target site by deleting, inserting, or replacing specific DNA sequences provides isogenic control cells to study pathogenic variants^5^.

Point mutations represent 58% of disease-related polymorphisms registered in the ClinVar database^6^. Precision gene editing technologies are required to reproduce these disease mutations in iPS cells at single nucleotide resolution^7^. The CRISPR-Cas9 system is conventionally used to generate targeted DNA double-strand breaks (DSBs) in the genome, which are subsequently repaired by cellular DNA repair pathways^8,9^. Non-homologous end-joining (NHEJ) results in insertion and deletion (indel) mutations, while microhomology-mediated end-joining (MMEJ) leads to predictable deletions^10^. Both NHEJ and MMEJ are collectively known as mutagenic end-joining (MutEJ), as both repair outcomes lead to the loss or gain of DNA sequence^11^. Typically, to produce specific changes including single nucleotide variants, the homology-directed repair (HDR) pathway is exploited with custom repair templates containing the desired edit, such as double-stranded linear or plasmid DNA, or single-stranded donor oligonucleotides (ssODNs). However, various studies have shown a preference for MutEJ over precise repair by HDR when employing canonical Cas9^12–15^.

Variations on Cas9, such as base editing (BE), combine Cas9 nickase (D10A) with a cytidine or adenine deaminase to directly convert specific DNA bases with reduced incidence of double-strand breaks^6,16^. Although BE can generate precise edits, it is limited to a specific editing window and bases adjacent to the target may be simultaneously converted, resulting in bystander mutations^6^. In contrast, Prime Editing (PE) technology combines Cas9 nickase (H840A) with reverse transcriptase (RT) activity derived from the Moloney murine leukemia virus (M-MLV)^17^. PE utilizes a prime editing guide RNA (pegRNA) with a 3’ extension that serves as a reverse transcription template (RTT) and primer binding site (PBS) to incorporate the desired edit. Since prime editing performs everything from single-stranded DNA cleavage to re-writing the genome, editing may be more intentional, and is a breakthrough in genetic disease modeling^18^.

This work demonstrates the optimization of PE applications in iPS cells using a GMP-grade electroporation platform. We established a fluorescence-based PE benchmark method by using GFP-to-BFP conversion in iPS cells. To maximize PE efficiency, we developed a strategy of FACS enrichment, in which the PE expression vector was modified with T2A-mCherry, allowing assessment of Prime Editor 2 (PE2) expression levels in cells and their fractionation. The efficiency of our method was benchmarked in iPS cells using *HEK3* and the rs429358 (c.T388C) pathogenic risk variant in *APOE*. Our results demonstrate that the activity of PE increases with the expression level of mCherry, and PE is less mutagenic than the conventional genome-editing method using Cas9. Moreover, our results demonstrate that FACS enrichment can be used to control allelic editing outcomes.

## Methods

### Human iPS cell culture

409B2 (RIKENBRC #HPS0076), 317-A4 (GFP heterozygously targeted iPS cells), and 317-D6 (GFP homozygously targeted iPS cells)^11,19^ were maintained at 37 °C and 5% CO2 in StemFit AK02N medium (Ajinomoto, Cat. No. RCAK02N) on 0.5 mg/mL silk laminin iMatrix-511 (Nippi, Cat. No. 892021) coated 6-well plates or 10 cm dishes. Cell passaging was performed every 7 days during maintenance. The cells were treated with 300 µL or 2 mL of Accumax (Innovative Cell Technologies, Cat. No. AM105-500) in 6-well plates and 10 cm dishes, respectively. The cells were incubated for 10 min at 37 °C to dissociate the cells. Pipetting was performed to detach the cells from the surface and generate a single-cell suspension in 700 µL or 4 mL of medium containing 10 µM ROCK inhibitor, Y-27632 (Wako, Cat. No. 253-00513). Cells were seeded onto iMatrix511-coated plates at a density of 1 × 10^3^ cells/cm^2^ in StemFit AK02N medium with 10 µM ROCK inhibitor for 24 h after seeding and then cultured without ROCK inhibitor. All the cell lines were routinely tested for mycoplasma contamination.

### Cas9-gRNA vector cloning

The spacer sequence of GFP-targeting gRNA was designed based on a previous work ^11^. pSpCas9(BB)-2A-GFP (PX458) was a gift from Feng Zhang (Addgene plasmid # 48138; http://n2t.net/addgene:48138; RRID:Addgene_48138). PX458 was digested by *Eco*RI, and T2A-mCherry was inserted and ligated (KW1013: pSpCas9(BB)-2A-mCherry). The gRNA construct was generated by Golden Gate assembly of annealed oligonucleotides into *the Bbs*I-digested KW1013 plasmid. The oligonucleotides listed in Table 2 were used for gRNA cloning.

### Cloning of PE-mCherry constructs

The T2A-mCherry fragment was PCR-amplified from the KW1013 plasmid. pCMV-PE2, a gift from David Liu (Addgene plasmid # 132775; http://n2t.net/addgene:132775; RRID; Addgene_132775), was digested by *Eco*RI and *Pme*I (Thermo Fisher Scientific). The digested product was gel-extracted using the Wizard SV Gel and PCR Clean-Up Kit (Promega). The two fragments were assembled in a single In-Fusion reaction (In-Fusion HD Cloning Kit, Takara, 639650), and the PCR-derived regions of the resulting plasmids were confirmed by sequencing.

### pegRNA design and cloning

pU6-pegRNA-GG-acceptor was a gift from David Liu (Addgene plasmid # 132777; http://n2t.net/addgene:132777; RRID; Addgene_132777). pegRNA-GFP and pegRNA-APOE were designed using the PrimeDesign web platform version (https://drugthatgene.pinellolab.partners.org/)^20^. The pegRNA construct was generated by Golden Gate assembly of annealed oligonucleotides into the *Bsa*I-digested pU6-pegRNA-GG-acceptor plasmid and sequence verified, as previously reported^17^. The plasmids used in this study are listed in Table 1, and the oligos used for the construction of vectors are listed in Table 2.

**Table 1.** Plasmids used in this study.

| Purpose | Plasmid ID | Plasmids |
| --- | --- | --- |
| ssODN editing | KW1013 | pSpCas9(BB)-2A-mCherry |
| ssODN editing | KW1322 | pSpCas9(BB)-2A-mCherry-GFP |
| PE | KW1564 | pCMV-PE2-T2A-mCherry |
| PE | KW1548 | pU6-pegRNA-GFP |
| PE | Addgene#132778 | U6-pegRNA-HEK3-ins3 (Anzalone *et al.*, 2019) |
| PE | KW1573 | pU6-pegRNA-GG-ApoE |

**Table 2.**
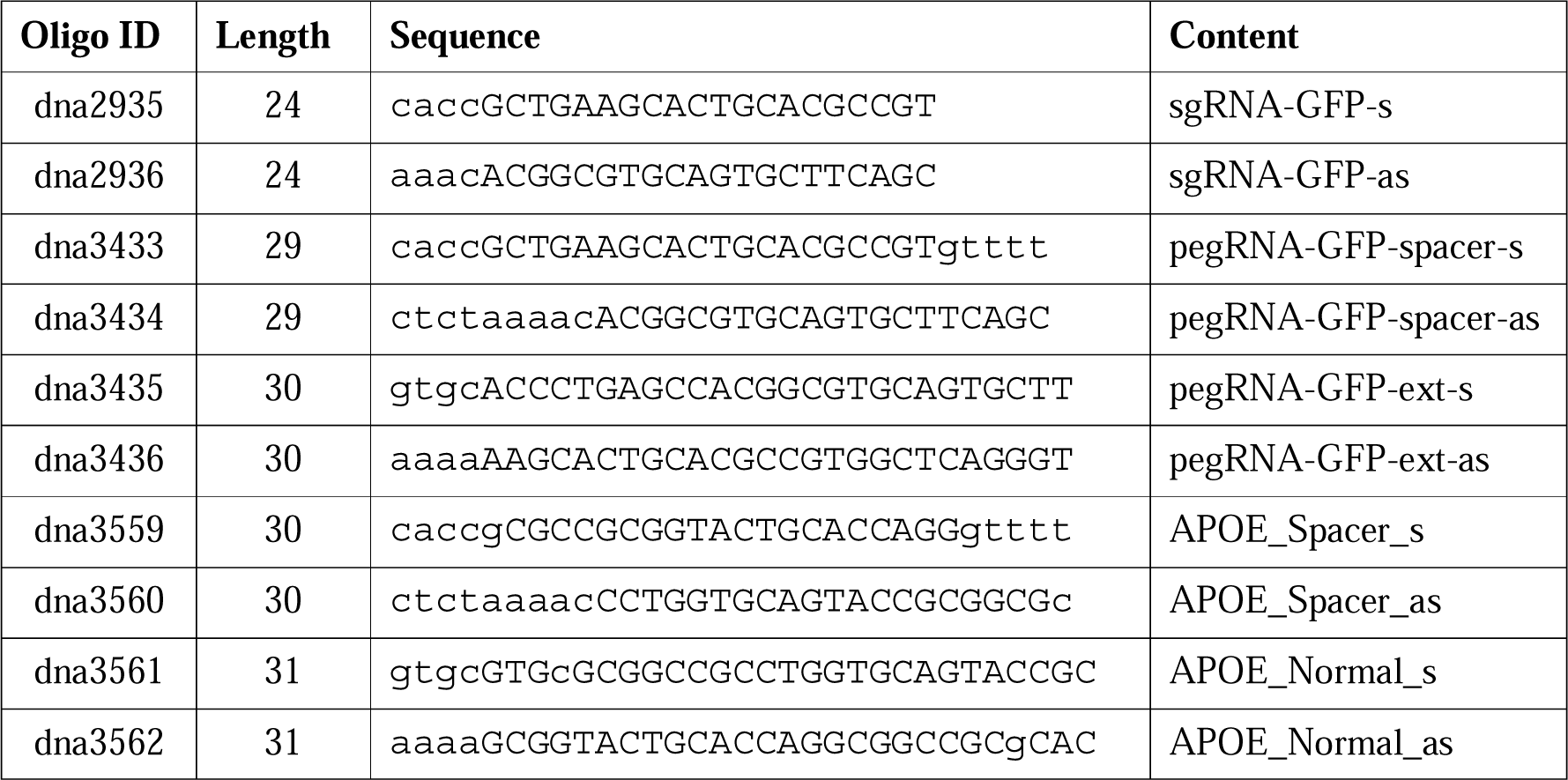
Oligos used for vector construction in this study.

### Electroporation of plasmids and RNP

For Cas9-based editing, all plasmids for electroporation were prepared using the HiSpeed Plasmid Maxi Kit (Cat. No. 12663), precipitated with ethanol, and dissolved in the MaxCyte electroporation buffer at a concentration of 2.5 µg/µL. For Cas9-based editing, 5 µg of plasmid encoding gRNA-GFP and Cas9 and 5 µg of ssODN repair template (IDT) were mixed in a total volume of 5 µL. The ssODNs used in this study are presented in Table 1. For PE-based editing, 5 µg of the KW1564 vector and 5 µg of the pegRNA-expressing plasmid were mixed in a total volume of 5 µL. Next, 5 × 10^6^ cells resuspended in 50 µL of MaxCyte electroporation buffer were added to the DNA mixture. The suspension (50 µL) was electroporated into an OC-100 × 2 processing assembly (MaxCyte, Cat. No. SOC-1 × 2) using a MaxCyte STX electroporator (Opt0-5 protocol). Electroporated cells were incubated at 37°C for 30 min and then transferred to an iMatrix511-coated 10 cm dish in StemFit AK02N medium supplemented with 10 µM ROCK inhibitor. Cell preparation for FACS analysis was performed 24 h after electroporation. Otherwise, medium exchange was performed 48 h after electroporation using StemFit AK02N without ROCK inhibitor, and cells were maintained until collection for genotyping on day five or flow cytometry (FC) analysis on day eight. We performed the *APOE* gene editing by ribonucleoprotein (RNP) of Cas9 with NEPA21 following the previously published protocol^11^. APOE gene editing by RNP was performed in T8 iPS cells^21^.

### Flow cytometry and cell sorting

5×10^5^ cells were suspended in 1 mL FACS buffer (DPBS supplemented with 2% FBS), and GFP and mCherry fluorescence intensities were analyzed using a BD LSRFortessa Cell Analyzer or BD FACSAria II cell sorter (BD Biosciences) with BD FACSDiva software (BD Biosciences). After setting gates for the singlets, 10,000 events were measured for each population. For editing experiments in 317-A4 iPS cells, the cells were acquired using Pacific Blue (450/50 nm) and FITC (530/30 nm) filters. For cell sorting, cell suspensions were prepared in FACS buffer at a density of 1 × 10^6^ cells/mL and filtered through a 35 µm nylon mesh cap of the tube (Corning, 352235) to remove clumps. Sorting gates were set for the singlet events. The desired population was collected using a BD FACSAria II cell sorter (BD Biosciences) in AK02N medium containing 20 µM Y-27632. Sorting efficiency was confirmed by re-analyzing 300 µL of the media. Raw data were analyzed using FlowJo 10 (FlowJo LLC). Rainbow Calibration Particles (6 peaks) and 3.0 - to 3.4 µm (BD biosciences) were utilized to calibrate the laser strength and determine the sorting gate. The 90 percentiles of the relative mCherry intensity of the unelectroporated cells was defined as the threshold of the mCherry.

### Genotyping

For genomic DNA extraction, 0.5–1 × 10^6^ cells were washed with 1X DPBS, and DNA was purified using the DNeasy Blood & Tissue Kit (Qiagen, Cat. No. 69506) following the manufacturer’s protocol. Purified DNA was eluted in 100 µL of AE buffer. Target sequences were amplified by PCR using the KAPA HiFi HS ReadyMix (Kapa Biosystems, Cat. No. KK2602). PCR product cleanup was performed using the ExoSAP-IT Express reagent (Cat. No. 75001) following the manufacturer’s protocol, and Sanger sequencing was performed using the BigDye Terminator v3.1 CS Kit (Thermo Fischer Scientific, Cat. No. 4337456). The final product was purified by ethanol precipitation and dissolved in HiDi formamide. Sequencing was performed on a 3500xl Genetic Analyzer (Applied Biosystems). Sequence alignments were analyzed with Snapgene (GSL Biotech LLC), and sequence trace files with low base-calling confidence were excluded manually. The primers used for genotyping are listed in Table 3. Sequencing analysis was performed on mixed sequences using ICE (https://ice.synthego.com/) and DECODR (https://decodr.org/) (REF)^22^. Sequence data from 317-A4 iPS cells was used as the reference genome. The parameters were kept at their default values.

**Table 3.**
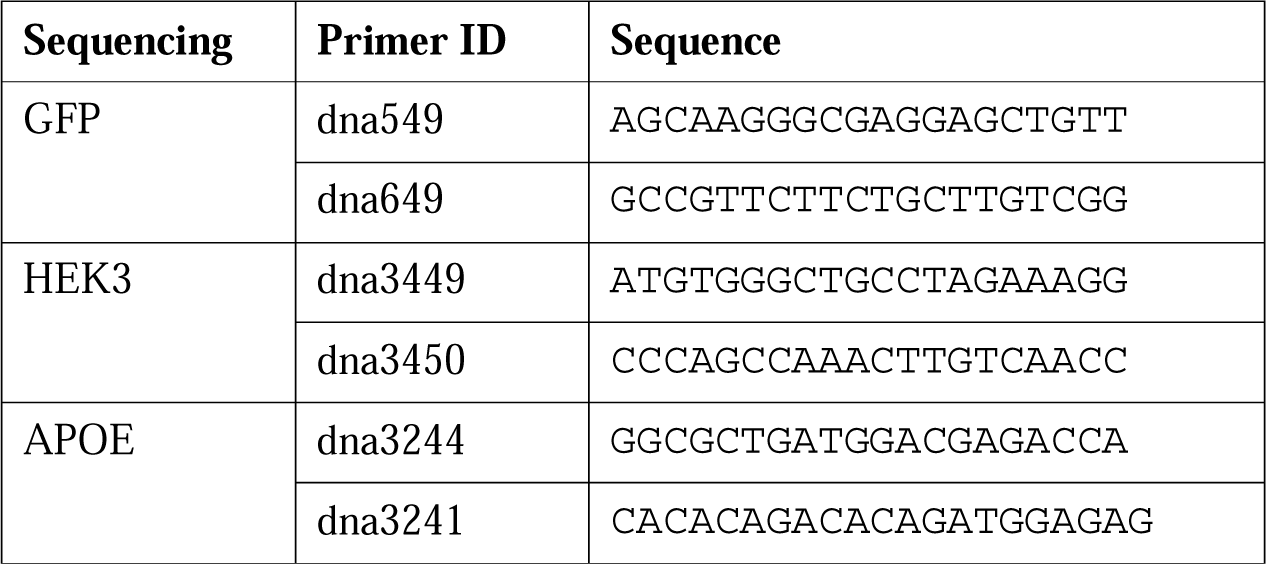
Primers used for genotyping and sequence analysis.

**Table 4.**
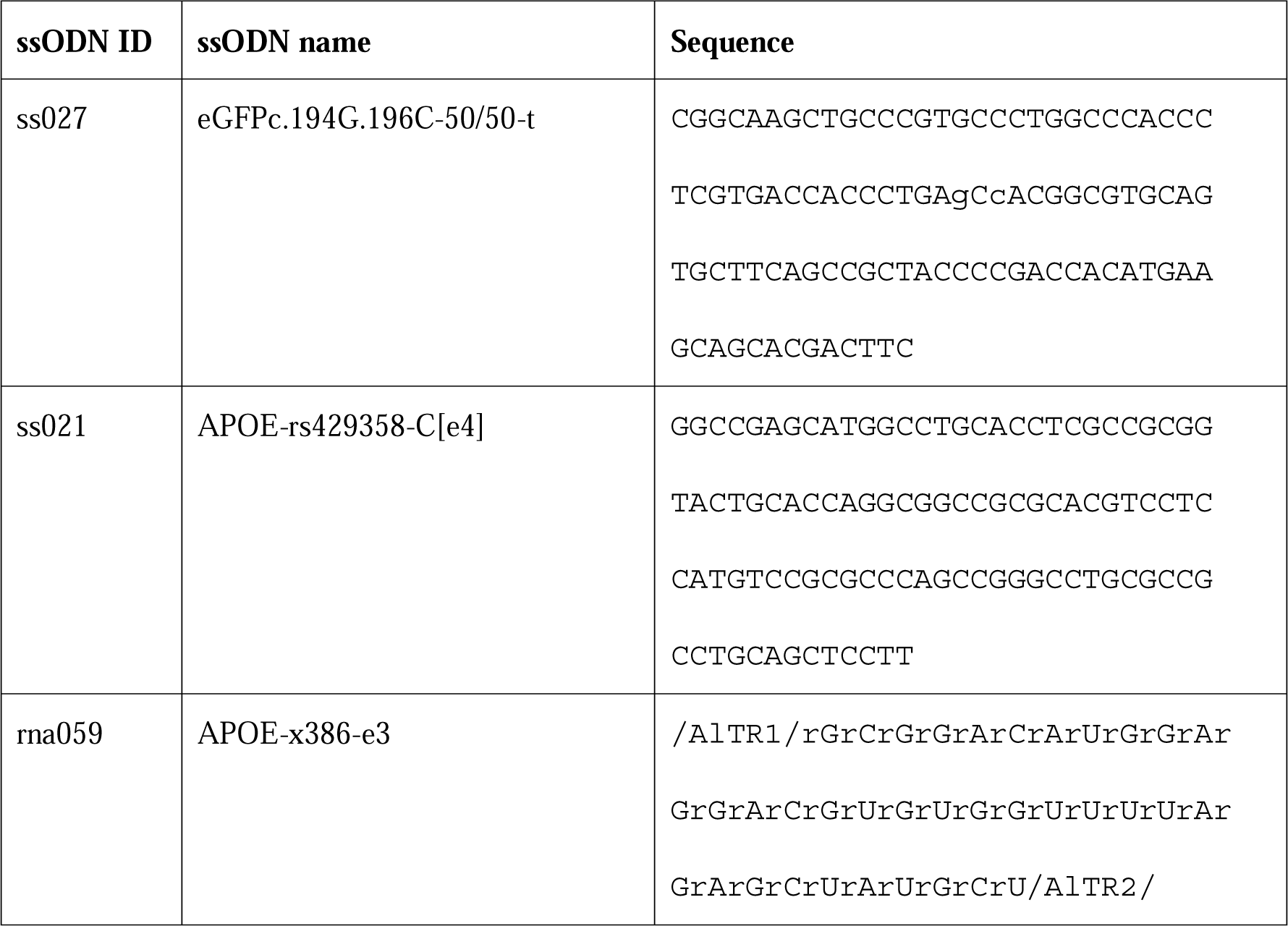
crRNA and ssODN used in this study.

### Droplet digital PCR (ddPCR)

To quantify the APOE 388 mutation created by PE, we prepared a mixture containing 10 µL of ddPCR Multiplex Supermix (Bio-Rad, Cat. No. 12005909), 1.8 µL of 20 µM forward and reverse primers each, 1.25 uL of 10 µM APOE-FAM and 0.5 µL of 10 µM hTFRC-HEX probes, and 30 ng of Template DNA, adjusted to a final volume of 20 µL. Droplets were generated using a QX200 Automated Droplet Generator (Bio-Rad). The PCR amplification protocol was as follows: 95°C for 10 min, 40 cycles at 94°C for 30 s and 58°C for 4 min, followed by 98°C for 10 min. The amplified droplets were then read with the QX200 Droplet Reader (Bio-Rad), and data were analyzed using QuantaSoft™ Analysis Pro v1.7.4.

### Statistical Analysis

The data are presented as the mean ± SD from the indicated numbers of independent experiments and were processed using R 4.0.3, and the R package tidyverse 1.2.0^23^ and ggprism^24^.

## Results

### GFP to BFP conversion assay to benchmark editing with PE plasmids in iPS cells

First, we established conditions for PE expression from plasmids in iPS cells. We adopted a GFP-to-BFP conversion assay previously used to optimize ssODN editing^11^, where a single amino acid change from tyrosine to histidine (Y66H) in the fluorophore region of GFP is sufficient to convert fluorescent emission of GFP to BFP, and can be quantified in single cells by flow cytometry (Figure 1a)^25^. The pegRNA-GFP was designed to create Y66H, along with a T65S mutation which acts to stabilize BFP, increasing fluorescence by approximately 2-fold^26,27^, as well as block the PAM to prevent subsequent re-cleavage of the edited BFP allele (Figure 1b). The spacer sequence is identical to that used in ssODN editing^11^. Conversion to BFP therefore represents the intended PE edit, while MutEJ is detected by loss of fluorescence and unmodified cells remain GFP-positive. These changes were quantified by flow cytometry, resulting in 21.2% editing to BFP in the 317-A4 iPS cell line (monoallelic AAVS1-targeted GFP) with 3 µg of each PE expression plasmid and pegRNAs expression plasmid (Figure 1c).

**Figure 1.**
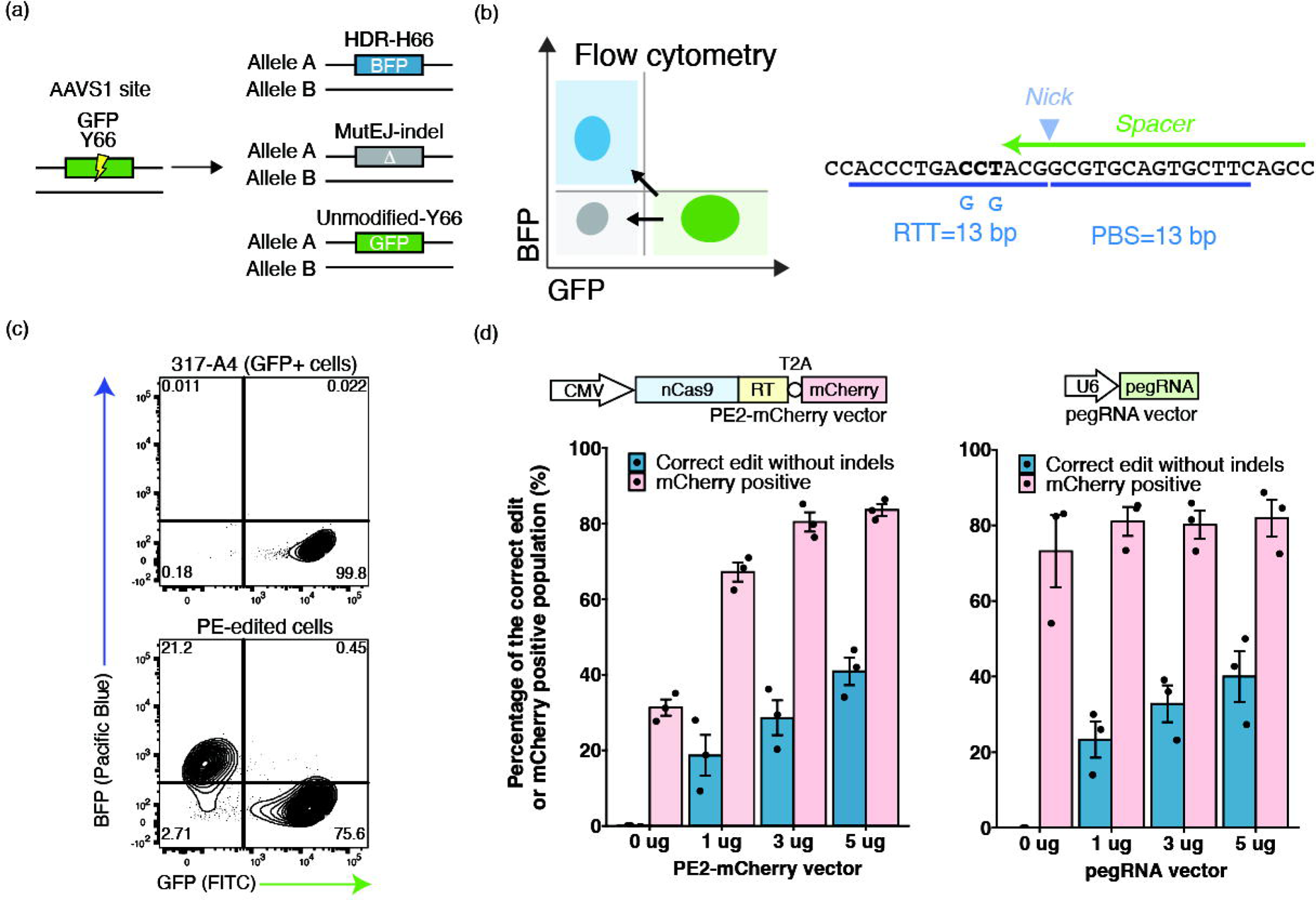
GFP-to-BFP conversion assay to optimize plasmid delivery of PE in human iPS cells using the MaxCyte platform. (a) Schematic overview of editing outcomes in heterozygous GFP reporter iPS cells (317-A4). A schematic of the Flow Cytometry (FC) distribution plot indicates each population. (b) Design of the pegRNA-GFP used in this study. The spacer sequence is indicated by a green arrow and the nicking position is indicated by the triangle. ssODN editing with Cas9 uses the same protospacer sequence^11^. RTT and PBS sequences are indicated by blue underlines. The PAM is in bold. Nucleotide changes leading to BFP are indicated in blue characters. (c) Representative plot of FACS analysis after GFP-to-BFP conversion. (d) Titration of PE2-mCherry (left) and pegRNA-GFP (right) plasmids. ‘mCherry positive’ indicates the proportion of cells over the 90 percentile of mCherry intensity in non-transfected cells at 24hrs after electroporation. ‘Correct edit’ indicates the proportion of BFP-converted cells on day 7 after electroporation.

The PE2 expression vector was modified to couple PE with mCherry using a T2A self-cleaving peptide (PE2-mCherry), such that mCherry expression represents the level of PE expression. Previously, delivery of Cas9 ribonucleoprotein (RNP) and DNA plasmid to human iPS cells was established on the GMP-compliant MaxCyte platform^11,28,29^. However, co-transfection of multiple plasmids in iPS cells has yet to be demonstrated, and we started by optimizing the amount of plasmids by titration. (Figure 1d). We tested the editing efficiency based on the amount of PE-mCherry or pegRNA expressing plasmid (0, 1, 3, and 5 µg), while keeping the second component fixed (5 µg). The proportion of mCherry-positive cells, detected at 24 h after electroporation, indicated transfection efficiency. The proportion of BFP-positive cells on day 7 indicated the amount of correct editing. We confirmed that the increase in mCherry-positive cells correlated with the amount of PE plasmid and the editing efficiency correlated with the amount of pegRNAs plasmid within the titration range. In addition, 5 µg of each plasmid showed consistent transfection efficiency of approximately 83% across multiple electroporations, and we decided to use this condition for the subsequent experiments. Notably, electroporation itself increased the autofluorescence and 31.3% was distinguished as positive when using 90 percentile of electroporated cells as a threshold.

### FACS enrichment of highly transfected iPS cells to maximize correct editing efficiencies

Next, we established parameters for FACS enrichment of cells expressing high levels of PE or Cas9. FACS enrichment was performed 24 hours after electroporation. Using fluorescent beads as a calibration ladder for consistency between experiments, the cells were divided into three groups: ‘Low’, ‘Medium’ (or ‘Med’), and ‘High’, depending on the level of mCherry expression (Figure 2a). The 90 percentiles of relative mCherry intensity of un-electroporated cells were used as a threshold for the Low population. The mode of each peak of the ladder was first measured, and we defined the mode of the 3^rd^ and 4^th^ peaks as the thresholds for Medium and High populations, respectively. The mCherry expression level varied among the population of transfected iPS cells (Figure 2b, c).

**Figure 2.**
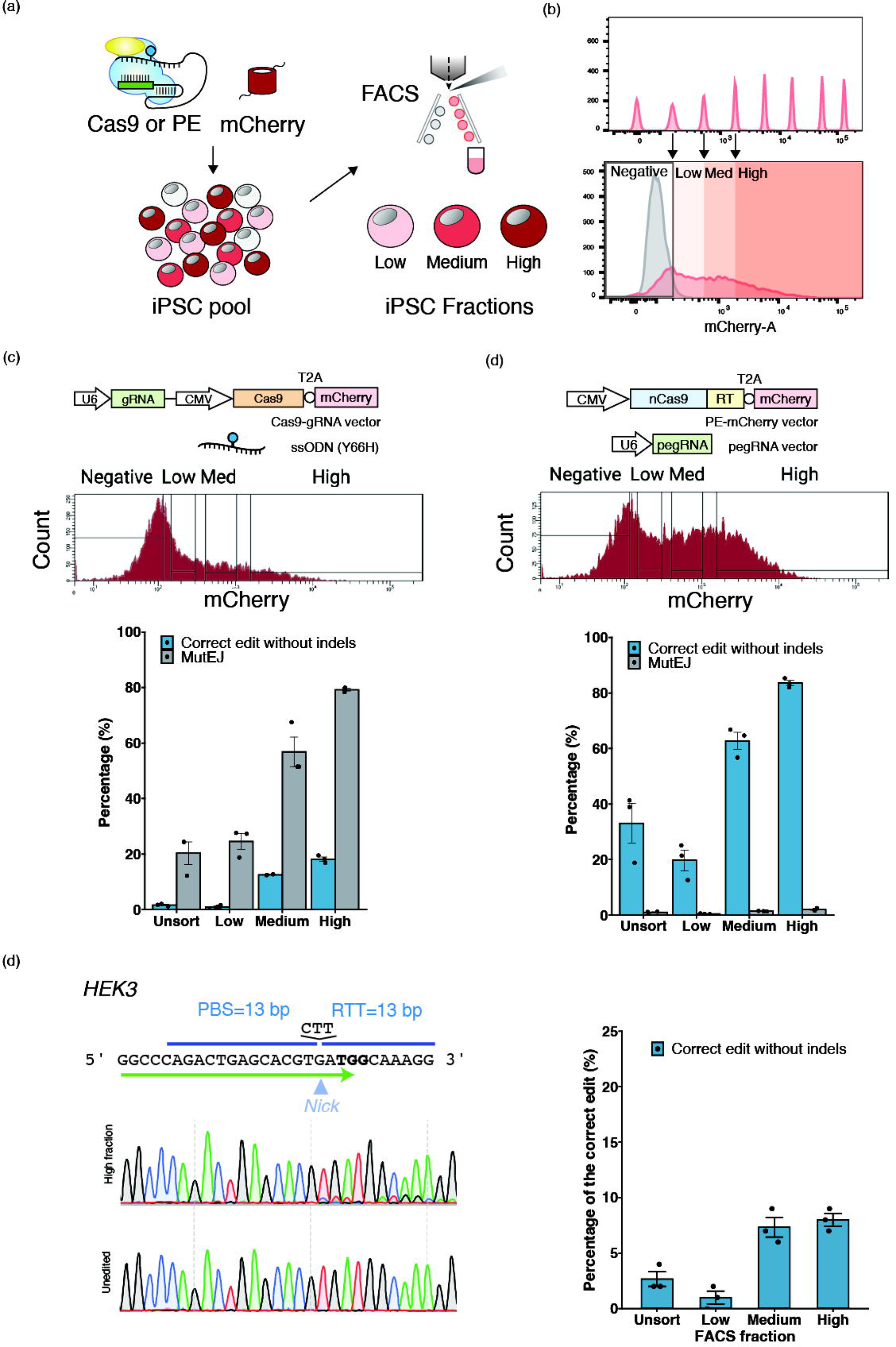
FACS enrichment to maximize PE or ssODN editing efficiency. (a) Schematic of FACS approach to cell fractionation based on transfection and expression level. Gating scheme was derived from a fluorescent calibration bead ladder and a representative histogram of mCherry positive cells with sorting gates. (c) Gating scheme and quantification of editing outcomes using Cas9-mCherry and ssODN-GFP. (d) Gating scheme and quantification of editing outcomes using PE and pegRNA-GFP. All data in (c, d) are presented as the meanl±lS.D. of three technical replicates of independent electroporations for each condition. (e) Design of pegRNA-HEK3 and editing outcomes as measured by Sanger sequencing.

The effect of FACS enrichment on the editing outcomes was tested for both PE and ssODN editing (Figure 2c, d). In the total unsorted (‘Unsort’) population, 33.0% of correct edits were observed for PE, whereas only 1.62% of the correct edits were observed in ssODN editing. MutEJ levels in PE were found to be low (0.993%), while MutEJ was more prevalent than correct edits for ssODN editing (20.3%). In comparison, the ratio of correct edits to MutEJ was 0.08 in ssODN editing, as compared to 33.3 in PE. These data verify that PE with plasmids is more efficient and precise than editing with Cas9 and ssODNs in human iPS cells. In the High fraction, on average, 83.6% of iPS cells were converted to BFP by PE while only 18.1% became BFP positive by ssODN. Compared with Unsort, the fold improvement of correct edits by PE were 0.60, 1.90 and 2.50 times in Low, Med, and High fractions, respectively. The fold improvement by ssODN editing was 0.53, 7.74, and 11.2 times for the respective fractions. Thus, for both PE and ssODN editing, the editing efficiency improved alongside mCherry intensity. Importantly for PE, the proportion of MutEJ only increased 0.43, 1.39, and 1.99 times in the Low, Medium, and High populations compared to Unsort, while in ssODN, MutEJ was increased 1.21, 2.80, and 3.90 times. In the High fractions, MutEJ reached only 18.1% for PE, but 79.2% for ssODN. These data demonstrate that FACS enrichment of PE increases the number of correct edits without a substantial increase in MutEJ, in stark contrast to ssODN editing. These data indicate that PE outcomes may be improved by enrichment using fluorescence, without an increase in MutEJ.

### Benchmarking editing efficiencies in iPS cells at endogenous loci

For benchmarking endogenous gene editing with PE and FACS enrichment, we selected the *HEK3* locus (Figure 2d) that has been used in diverse cell lines such as HEK293T, HeLa, K562, and human embryonic stem (ES) cells^30,31^. This benchmarking pegRNA inserts CTT and caused two detected sequences. Overall, the combination of PE and FACS enrichment resulted in a 3-fold increase in correct edits, reaching 8.0% in the High fraction. MutEJ was undetectable across all fractions by ICE analysis. These data indicate that PE can be improved by fluorescent enrichment at endogenous loci, without co-enrichment of MutEJ.

### Evaluation of allelic editing outcomes with FACS enrichment

Given the potential of iPS cells for modeling genetic diseases, it is important to determine the rate of mono- and biallelic editing at a cellular or clonal population level. We first evaluated allelic editing in 317-D6, a biallelic AAVS1-targeted GFP iPS cell line. In total, six editing patterns are expected (Figure 3a). Since iPS cells with one or two active copies of GFP or BFP only double in their mean fluorescence intensities^19^, we recognize that distinguishing between biallelic editing and monoallelic editing with MutEJ is challenging by FACS^11^. However, considering the near-zero proportion of MutEJ generated by PE, we predicted that BFP single-positive iPS cells correspond to biallelic editing, whereas BFP/GFP double-positive iPS cells represent monoallelic editing (Figure 3a). The Unsort population exhibited nearly equal amounts of biallelic (29.4%) and monoallelic (23.1%) editing (Figure 3b). Interestingly, the High population showed more than 3-fold higher biallelic editing (72.6%) than monoallelic editing (22.4%). In the Med population, these proportions were more similar at 43.1% and 36.3% for biallelic and monoallelic edits, respectively. Finally, the Low population reversed this trend, showing nearly half the number of biallelic edits compared to monoallelic editing (7.65% and 14.7%). These results demonstrate that FACS enrichment for defined PE expression levels can skew the outcomes of mono- and biallelic editing at a cellular level.

**Figure 3.**
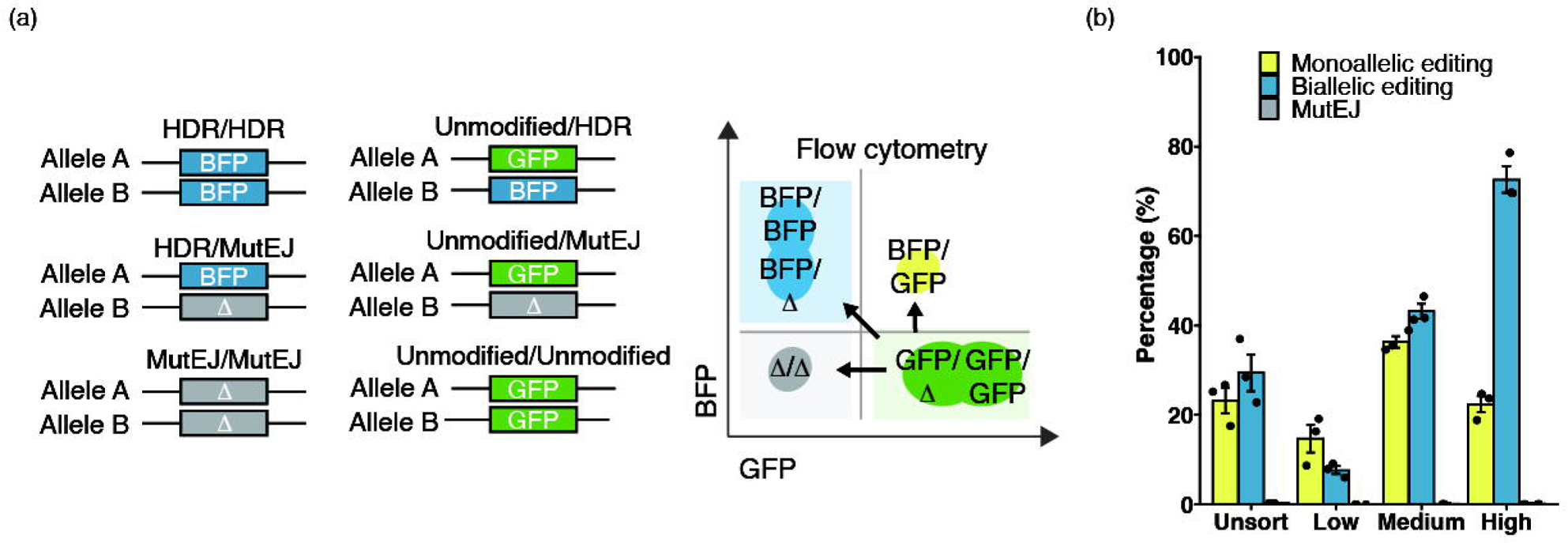
Modulating the allelic editing outcomes by FACS enrichment. (a) Schematic of the expected GFP editing outcomes in a homozygous AAVS1-CAG::GFP iPS cell line. (b) Allelic editing outcomes following FACS enrichment (N=3). Each fractioned population showed different pattern in editing outcomes.

We then explored the clonal distribution of allelic editing by Cas9 and PE at an endogenous locus. Among the three variants of the apolipoprotein (*APOE*) gene, APOE2, APOE3 (c.C526T, rs7412), and APOE4 (c.T388C, rs429358), APOE4 is associated with the highest risk of Alzheimer’s disease (Figure 4a). We therefore chose to engineer the rs429358 APOE3 variant to the APOE4 genotype by both ssODN editing and PE (Figure 4b). Editing by ssODN and Cas9 RNP showed 11.3% correct edits (Figure 4c). However, correct edits were outcompeted by 78.3% +1T insertions. We treated cells with NU7441 (a DNA-PKcs inhibitor), with the aim of depleting +1T insertions and increasing the correct edits (Figure 4d). NU7441 treatment was able to maintain a similar level of correct edits (7.5%). While the decrease in +1T insertion by NU7441 was substantial (55.0%), it was not enough to eliminate it. Next, we created the same variant by PE and FACS enrichment (Figure 4e, f). Sanger sequencing with DECODR software estimated 23.6% APOE4 c.T388C correctly edited alleles in the High fraction, which was validated by droplet digital PCR (ddPCR, 21.3%) (Figure 4f). In contrast, no edits were detected in the Low or Med fractions by Sanger sequencing and DECODR, whereas ddPCR identified 3% and 8% correct edits, respectively (Figure 4f). MutEJ was not discernible in any of the fractions by Sanger sequencing analyzed by DECODR.

**Figure 4.**
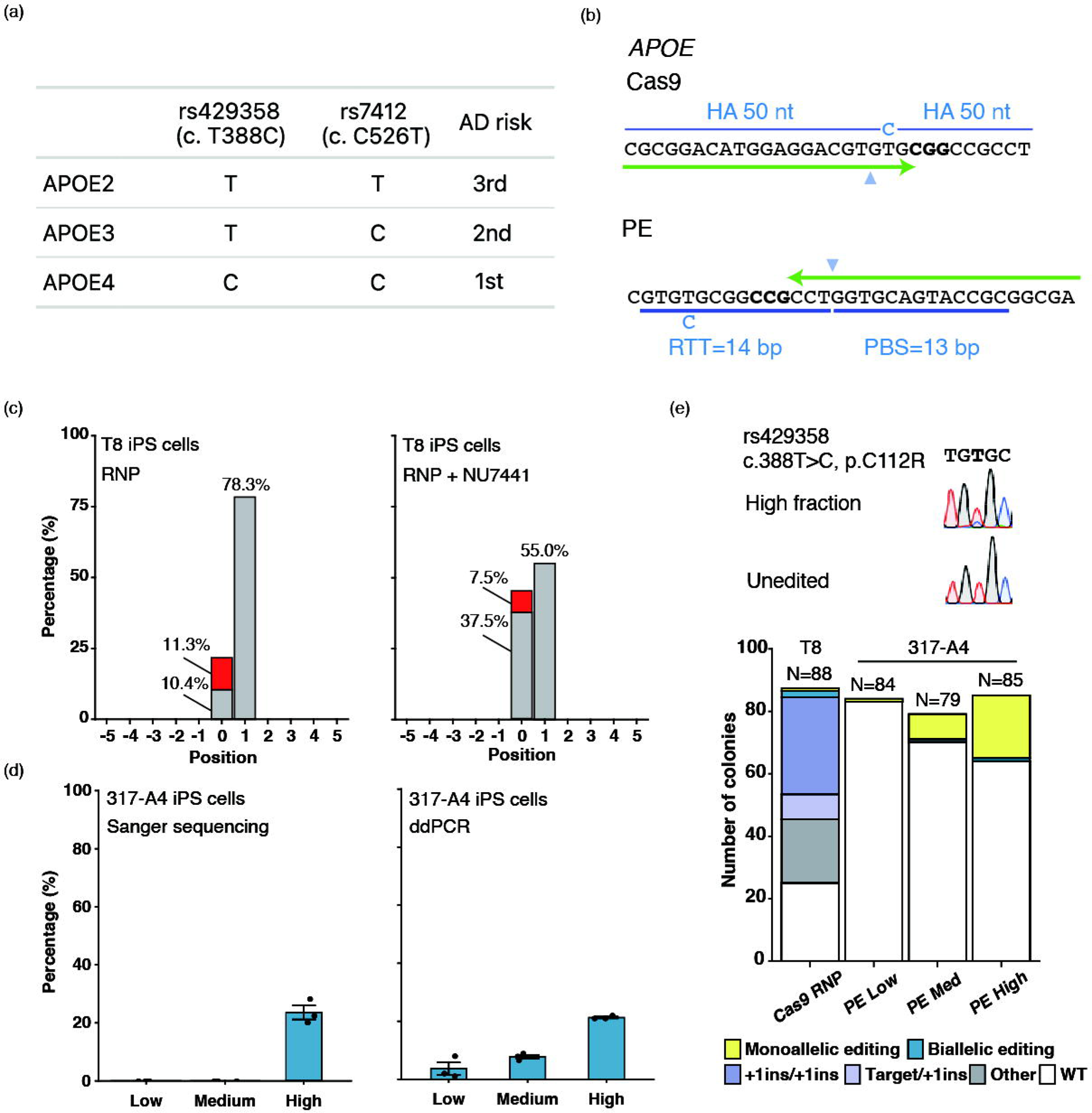
APOE gene editing to recreate a disease model for Alzheimer’s disease in iPS cells. (a) Table of APOE risk variant alleles for Alzheimer’s disease. (b) Schematic of the design for gRNA- and pegRNA-APOE. (c) TIDE plot of ssODN editing in T8 iPS cells. (d) ssODN editing with NU7441 to reduce +1 insertion. (e, f) Quantification of Correct edits by DECODR analysis (e) and ddPCR (f) in 317-A4 iPS cells after PE and mCherry FACS. (g) Clonal distribution of gene editing outcomes at the APOE locus using ssODN or PE.

Finally, we determined the number of APOE4 biallelic and monoallelic edited iPS cells at a clonal level (Figure 4g). Using Cas9 RNP, two bi- and nine monoallelically edited clones were obtained, however, 8 of the monoallelically edited clones were accompanied by a +1 insertion in other alleles. Using PE, the Med and Low populations yielded only monoallelic clones (1 and 8, respectively), with many unedited clones. From the High PE condition, we isolated twenty-four mono- and one biallelically edited colony. Moreover, no unintended edits were confirmed by Sanger sequencing of clones. Collectively, these findings suggest that a combination of PE and FACS enrichment can fulfill a crucial need in establishing an allelic series of isogenic iPS cells for disease modeling.

## Discussion

In this research, we demonstrate the superior efficiency and precision of PE over Cas9-based ssODN editing in iPS cells. Also, applying FACS, we successfully enriched cells exhibiting varied editing efficiencies. Leveraging a GFP to BFP reporter assay with single-cell resolution, our findings revealed that PE generated notably fewer MutEJ events compared to ssODN editing. Finally, we show that FACS enrichment can modulate the frequency of mono- or bi-allelic editing, demonstrating its utility in controlling allelic editing outcomes.

The GMP-approved MaxCyte platform has been used for plasmid transfection into T cells, HEK293T cells, and CHO cells^32,33^. Previous research used the MaxCyte to successfully introduce plasmid, mRNA, and RNPs into iPS cells^11,28,29,34^. As iPS cells hold significant promise not only for disease modeling but also for genome-edited cellular regenerative medicine anticipating ex vivo applications for cell therapy, our findings are of paramount importance.

Our protocol demonstrates a highly reproducible FACS gating strategy through the implementation of a fluorescent bead ladder. FACS provides advantages for PE editing using plasmid vectors by eliminating non-transfected cells and isolating cells based on their expression level. Our FACS enrichment data also indicate that PE activity was underestimated in the unsorted population, similar to a previous study on the combination of Cas9 and FACS enrichment reported^35^. Another study employed *piggyBac* transposition to introduce PE into iPS cells for long-term stable expression, yet observed no increase in editing over 40 days of continuous expression^36^, suggesting that the effects of PE are realized early after transfection. Using FACS enrichment, it is possible to select cells with modest PE expression for editing with high efficiency pegRNAs (such as pegRNA-GFP), and high PE expression for editing with low efficiency pegRNAs (such as pegRNA-APOE).

To bias the reaction of heteroduplex removal in the PE repair step, a second nicking guide RNA (ngRNA) can be used to nick approximately 50 bases in the editing region. Prime editing with the ngRNA used is called PE3^16,17,37^. Although previous studies have shown that PE3 can be further improved by additional nicking, this also increases MutEJ. Also, using multiple gRNA including pegRNA in a single experiment increases the sites experimentalists require to confirm off-target activity. Thus, our study focused on PE2. Editing of GFP was found to be efficient; however, editing at endogenous loci was variable. PE3 can be a powerful method for improving editing efficiency. Furthermore, in PE, almost all edits were targeted edits. Previous studies have shown that PE is less likely to occur in cells with mature Mismatch Repair (MMR) such as iPS cells^30^. PE4 is a method in which the transient expression of a dominant-negative MMR protein (MLH1dn) is combined with PE2. This temporal inhibition of MMR showed the potential to increase editing efficiency in iPSCs^38^. Combined with FACS enrichment, this method may enable further flexible and accurate genome editing.

Even without FACS, we observed a significant disparity in by-products between ssODN editing and PE. Upon FACS enrichment, ssODN editing showed a marked increase in the proportion of MutEJ accompanied by HDR, whereas, in PE, only the proportion of correct edits surged. This disparity is caused by the increased likelihood of DSBs due to elevated Cas9 expression levels, whereas PE’s nick-based editing mechanism enhances mainly the rate of correct editing over MutEJ. While our observations were consistent for editing a single nucleotide variant in the *APOE* gene, we cannot guarantee that similar patterns would be observed for other genes or pegRNA and mutation designs without a genome-wide effort. In addition, pegRNA activity differs between *GFP* and *APOE*. This resulted in high Bi-allelic editing in *GFP*, whereas high Mono-allelic editing was observed in *APOE*. Allelic editing may require a high level of PE activity.

## Conclusion

Combination of PE and FACS enrichment allows highly efficient and accurate gene editing. The combined protocol is a useful method, as the accuracy of PE does not lead to a significant increase in by-products. Also, fractionation by FACS based on PE expression level can influence mono- or bi-allelic editing. Improving the reliability of gene editing outcomes will greatly facilitate the generation of genetic disease models and possibly therapies using human iPS cells.

## Acknowledgment

We thank Suji Lee for technical assistance in cell culture and molecular biology, Dr. Kanae Mitsunaga for helping establish conditions for FACS analysis and cell sorting, and Dr. Peter Gee for helping establish conditions for large plasmid delivery on the MaxCyte and reviewing the manuscript. We thank Dr. Ryo Yamada for his mentorship and guidance of R.N. as part of the Medical Innovation Program and providing the inspiration for statistical methods applied to FACS.

## Funding information

R.N. is a trainee of the Medical Innovation Program managed by Kyoto University and supported by JST SPRING (Grant Number JPMJSP2110). This study was supported by funding to K.W. from the Japan Agency for Medical Research and Development (AMED) for the Core Center for iPS Cell Research (Grant Number JP21bm0104001), the Core Center for Regenerative Medicine and Cell and Gene Therapy (Grant Number JP23bm1323001), the COVID-19 Private Fund (Shinya Yamanaka, CiRA, Kyoto University), and the Canadian National Research Council.

## Competing Interest Statement

The authors declare that they have no conflicts of interest.

## Author contribution

RN: Conceptualization, Formal analysis, Investigation, Resources, Writing - Original Draft, Writing - Review & Editing, Visualization

TM: Methodology, Investigation, Resources

AYL: Investigation, Resources

MK: Investigation

TK: Investigation, Resources KT: Resources

HI: Investigation, Resources

TLM: Conceptualization, Investigation, Writing - Review & Editing

KW: Conceptualization, Formal analysis, Investigation, Resources, Writing - Original Draft, Writing - Review & Editing, Supervision, Project administration, Funding acquisition

## Notes

### Competing Interest Statement

The authors have declared no competing interest.

